# Dynamical Model of Drug Accumulation in Bacteria: Sensitivity Analysis and Experimentally Testable Predictions

**DOI:** 10.1101/030908

**Authors:** Neda Vesselinova, Boian S. Alexandrov, Michael E. Wall

**Affiliations:** Theoretical Division, Los Alamos National Laboratory, Los Alamos NM 87545; University of California, Los Angeles, CA 90095; Computer, Computational, and Statistical Sciences Division, Los Alamos National Laboratory, Los Alamos NM 87545

## Abstract

We present a dynamical model of drug accumulation in bacteria. The model captures key features in experimental time courses on ofloxacin accumulation: initial uptake; two-phase response; and long-term acclimation. In combination with experimental data, the model provides estimates of import and export rates in each phase, the time of entry into the second phase, and the decrease of internal drug during acclimation. Global sensitivity analysis, local sensitivity analysis, and Bayesian sensitivity analysis of the model provide information about the robustness of these estimates, and about the relative importance of different parameters in determining the features of the accumulation time courses in three different bacterial species: *Escherichia coli, Staphylococcus aureus*, and *Pseudomonas aeruginosa*. The results lead to experimentally testable predictions of the effects of membrane permeability, drug efflux and trapping (e.g., by DNA binding) on drug accumulation. A key prediction is that a sudden increase in ofloxacin accumulation in both *E. coli* and *S. aureus* is accompanied by a decrease in membrane permeability.

**Author Summary:** Bacteria live or die depending on how much antibiotic gets inside them. Using a simple mathematical model, detailed information about drug import and export can be teased out of time courses of internal drug levels after a sudden exposure. The results suggest that membrane permeability can suddenly decrease during exposure to drug, accompanied by an increase, rather than a decrease, in the internal drug level.

## Introduction

Drug resistance in bacteria can be increased by efflux pump systems [1], and pump inhibition has emerged as a strategy for overcoming drug resistance [2]. Many details of how efflux pumps work are still unclear, however. In particular, quantitative information about how efflux influences drug accumulation in bacteria is still scarce [3].

Drug accumulation is a key factor in obtaining a quantitative understanding of resistance. For example, predictions of minimum inhibitory concentrations (MICs) of β-lactams in *Escherichia coli* were obtained by equating the steady state periplasmic drug concentration with a periplasmic binding protein inhibitory concentration [4]. A predicted MIC was calculated as the *external* concentration that would yield the accumulated *internal* inhibitory concentration in steady state, considering flux terms from membrane permeation and β-lactamase degradation. MIC predictions also have been made considering the action of efflux pumps on cephalosporins [5] and β-lactams [6] in *E. coli*. These predictions were accompanied by estimates of efflux pump kinetic constants (*i.e*., *K_m_* and *V_max_* values), providing an explicit connection between efflux and resistance.

Time-dependent drug accumulation studies also have yielded insight into drug transport [7–9]. Diver et al. [8] found exposure of *E. coli* to five different quinolones induced a rapid ̴10 sec uptake followed by a ̴30 min phase of more gradual 3 accumulation. Similar two phase behavior was seen by Asuquo and Piddock [7] in accumulation of fifteen different quinolones in *E. coli, Pseudomonas aeruginosa*, and *Staphylococcus aureus* [7]. The experiments of Asuquo and Piddock [7] highlighted exposure to ofloxacin, which exhibited apparently different accumulation behavior in each species (Fig. 1). Whereas drug levels appeared to plateau in *E. coli* (Fig. 1A) and *S. aureus* (Fig. 1C) levels in *P. aeruginosa* gradually decreased at longer times (Fig. 1B), suggesting an acclimation process.

**Figure 1.**
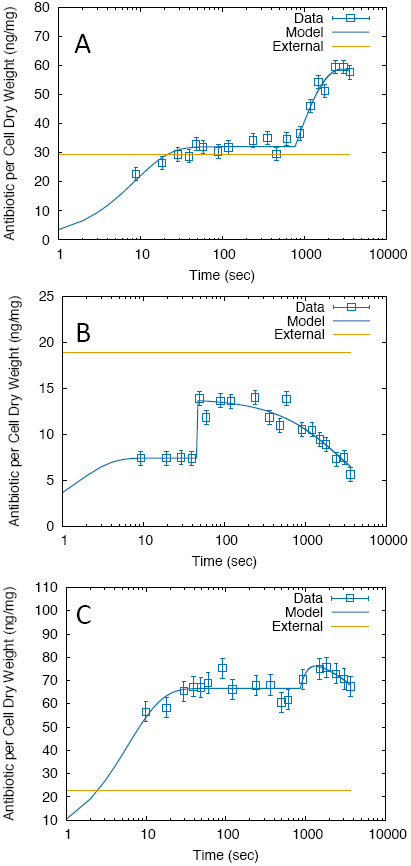
Ofloxacin accumulation data (open squares) with model fits (solid lines). A) *E. coli*. B) *P. aeruginosa*. C) *S. aureus*. Each data point is assumed to have the same measurement error, estimated from the maximum values reported for norfloxacin in Ref. [7] (vertical bars). The external drug concentration of 10 mg/L is shown in each panel in ng/mg units (dashed straight lines), converted using the factors derived from the buoyant densities and dry fraction (Methods). The x-axis is plotted using a log scale, reflecting the time sampling of the data.

Mathematical modeling has provided substantial insights into accumulation of β-lactams [4,6], cephalosporins [5], and tetracycline [9] in *E. coli*. Quinolone accumulation, however, has not yet been analyzed in the context of a mathematical model. Here we present a mathematical model of drug accumulation and use it to analyze experimental data on accumulation of the quinolone ofloxacin in *E. coli, S. aureus*, and *P. aeruginosa [7]*. The analysis yields estimates of permeation and efflux rates, the time of entry into the second phase of the response, and the rate of decrease of drug during acclimation. We also perform global sensitivity analysis, local sensitivity analysis, and Bayesian sensitivity analysis of the model. The sensitivity analyses provide information about the robustness of parameter estimates and enable assessment of the relative importance of the short-term response and long-term acclimation in different bacterial species.

The results lead to experimentally testable predictions of membrane permeability and drug efflux, which influence drug resistance. A key prediction is that an increase in *E. coli* and *S. aureus* drug accumulation is accompanied by a decrease in membrane permeability. Overall the results indicate the utility of mathematical modeling and sensitivity analysis in obtaining new insights into drug accumulation in bacteria.

## Results

### Model behavior

Both the *E. coli* (Fig. 1A) and *P. aeruginosa* (Fig. 1B) behavior clearly show two phases of drug accumulation dynamics, which is captured by our model of drug accumulation (Methods). The model also allows for the possibility of acclimation in the second phase; this was initially added to capture the longer-time (̴ 10-60 min) behavior of *P. aeruginosa* (Fig. 1b). Although *S. aureus* data initially appeared to involve only one phase (Fig. 1C), the modeling and sensitivity analysis later revealed a significant two-phase behavior, with acclimation.

The relationship between the model parameters and the model behavior is illustrated in Fig. 2. After exposure at *t* = 0, the first phase begins, where cellular antibiotic levels rise and relax toward an asymptotic value *a*_1_. The relaxation in this phase is exponential with rate β_1_. At time *t* = τ, the second phase begins, where antibiotic levels may either rise or fall towards a new asymptotic value *a*_2_. The relaxation in this phase is initially dominated by exponential relaxation with rate β_2_.

**Figure 2.**
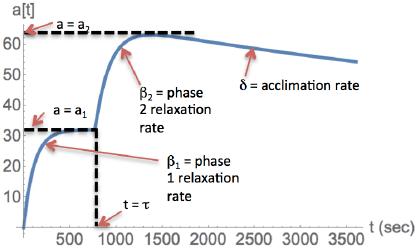
Behavior of the model and connection to parameters.

Later, acclimation can dominate the dynamics, specified by modulation of β_2_ by the factor [1+δ(t-τ)] (or, equivalently, modulation of α_2_ by [1+δ(t−τ)]^−1^; see Discussion). Together these features capture the full set of salient behaviors exhibited by the experimental data on ofloxacin accumulation [7] (Fig. 2).

### Parameter estimates

Optimal parameter values were estimated by fitting the model to data on ofloxacin accumulation in *E. coli*, *P. aeruginosa*, and *S. aureus*. The fits were performed assuming the same measurement error for each data point within the same curve (Methods). Reasonable fits were obtained for all datasets (Fig. 1), with an optimal value of the goodness of fit χ^2^/NDF of 0.98,1.5, and 0.74 for *E. coli*,*P. aeruginosa*, and *S. aureus*, respectively. Values of β_1_, β_2_, and τ for *E. coli* and *S. aureus* were surprisingly similar (Table 1) given the visible differences between the two datasets (Fig. 1); the parameter values for *P. aeruginosa* differed from the other two cases (Table 1). *E. coli* exhibits two phases of accumulation with little apparent long-term acclimation, consistent with the low value of δ. *P. aeruginosa* shows a stronger acclimation response, corresponding to the higher value of δ. *S. aureus* exhibits a degree of acclimation (δ) that is intermediate between *E. coli* and *P. aeruginosa*.

**Table 1.** Estimated parameter values obtained after fitting the model to data on ofloxacin accumulation in different bacteria.

| | $a_1$ | $a_2$ | $\beta_1$ (1/s) | $\beta_2$ (1/s) | $\delta$ (1/s) | $\tau$ (s) |
| --- | --- | --- | --- | --- | --- | --- |
|  | (ng/mg) | (ng/mg) |  |  |  |  |
| *E. coli* | 32.0 | 67.1 | 0.116 | 0.00125 | 0.000048 | 772 |
| *P. aeruginosa* | 7.43 | 13.7 | >0.1<sup>a</sup> | >0.1<sup>a</sup> | 0.000314 | 44.5 |
| *S. aureus* | 66.6 | 85.2 | 0.169 | 0.0014 | 0.000091 | 740 |

<sup>a</sup>Entries using '>' represent lower bounds.

### Sensitivity analysis results

We applied three types of sensitivity analysis to the model: global sensitivity analysis (GSA), local sensitivity analysis (LSA), and Bayesian sensitivity analysis (BSA). The GSA yielded a fine-grained assessment of the importance of each parameter in determining the model behavior at each time point. The LSA indicated parameters that are responsible for the principal variations in the immediate neighborhood of the optimal values. The BSA provided a detailed picture of the shapes of probability distributions of parameters in a more extended range about the optimum than for the LSA, including correlations that arise due to the nature of the model equations. Semi-quantitative sensitivity rankings of parameters for each case are presented in Supplementary Table S1; the detailed results are described below.

## Global Sensitivity Analysis (GSA)

GSA results were obtained using log-uniform prior distributions over the ranges in Table 2. The results for *E. coli* (Fig. 3A,B) and *S. aureus* (Fig. 3E,F) are similar, due to the identical priors (Table 2) and the similarity in the optimal parameter values for these cases (Table 1). The “main effect” index (a.k.a. the first-order sensitivity index) quantifies the contribution to the variance of the model of each parameter alone, while averaging over variations in other parameters, whereas the “total effect” index quantifies the change in the variance of the model caused by the investigated parameter and its interactions with any of the other parameters (Methods). Both the main effect and total effect are measured on a scale from 0 to 1 for each parameter.

**Figure 3.**
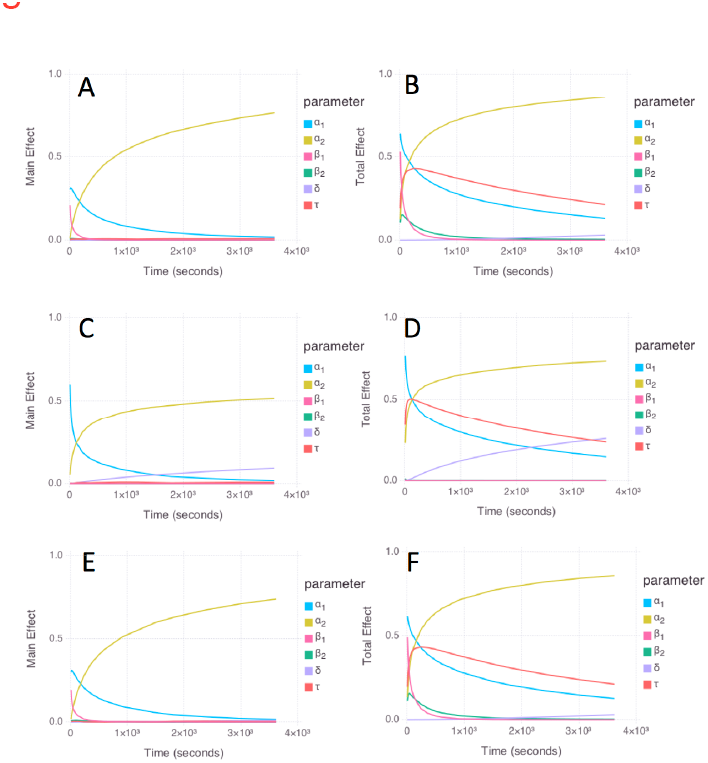
Global sensitivity analysis results for *E. coli* (A, B); *P. aeruginosa* (C, D); and *S. aureus* (E, F) models. The left panels (A, C, E) show the main effect results, and the right panels (B,D,F) show the total effect results. Color code: *a*_1_ (cyan); *a*_2_ (yellow); β_1_ (magenta); β2 (green); δ (lavender); and τ (red).

**Table 2.** Parameter ranges used for prior distributions for GSA and BSA (Log_10_ transformed values). Units of parameters are as in Table 1.

| | Log<sub>10</sub> $a_1$ | Log<sub>10</sub> $a_2$ | Log<sub>10</sub> $\beta_1$ | Log<sub>10</sub> $\beta_2$ | Log<sub>10</sub> $\delta$ | Log<sub>10</sub> $\tau$ |
| --- | --- | --- | --- | --- | --- | --- |
| *E. coli* |  |  |  |  |  |  |
| min | 0 | 0 | -3 | -3 | -7 | 0 |
| max | 2 | 2 | 0 | 0 | -3.5 | 4 |
| *P. aeruginosa* |  |  |  |  |  |  |
| min | 0 | 0 | -1 | -1 | -7 | 0 |
| max | 2 | 2 | 4 | 4 | -2 | 4 |
| *S. aureus* |  |  |  |  |  |  |
| min | 0 | 0 | -3 | -3 | -7 | 0 |
| max | 2 | 2 | 0 | 0 | -3.5 | 4 |

As a reference for the following paragraphs, *a*_1_ and *a*_2_ correspond to the initial steady-state level of antibiotic in the first and second phase; β_1_ and β_2_ correspond to the relaxation rates at the entry into the first and second phase; τ is the time of entry into phase 2; and δ is the rate constant associated with acclimation.

For *E. coli* and *S. aureus*, the main effect of *a*_1_, *a*_2_, and β_1_ is high, and the main effect of β_2_, τ, and δ is low (Figs. 3A,3E). The total effect of *a*_1_, β_1_, *a*_2_, and τ is high, and the total effect is low for β_2_ (especially at later times) and δ (Figs. 3B, 3F). The total effect of *a*_1_ remains high at long times, despite being associated with the first phase; this is because the prior for τ spans the entire range 0–3600 s, enabling the first phase to extend to long times in the analysis.

For *P. aeruginosa*, the main effect of *a*_1_, *a*_2_, and δ is high, and the main effect of β_1_, β_2_, and τ is low (Fig. 3C). The total effect of *a*_1_, *a*_2_, τ, and δ is high, and the total effect of β_1_ and β_2_ is low (Fig. 3D). The increased importance of δ for *P. aeruginosa* compared to *E. coli* and *S. aureus* is consistent with the more pronounced acclimation response (Fig. 1). The low importance of β_1_ and β_2_ is consistent with the unbounded upper limit due to the step-like transitions in *P. aeruginosa;* in comparison, the behavior of *E. coli* and *S. aureus* shows more gradual relaxations with improved sampling (Fig. 1), leading to increased importance of β_1_ and β_2_.

## Local sensitivity analysis (LSA)

The LSA results are presented using the eigenvectors and eigenvalues of the covariance matrix of model parameters (Methods). The eigenvectors are visualized in columns of colored squares in Fig. 4, ordered in importance from left to right using the eigenvalues (Table 3; sensitivity values correspond to −Log_10_(λ), where λ is the eigenvalue). For all cases, the first two eigenvectors are dominated by *a*_1_ and *a*_2_, indicating that these are the most sensitive parameters. For *E. coli*, the third eigenvalue is more than 10-fold larger than the first eigenvalue (corresponding to a lower sensitivity), with an eigenvector that is dominated by τ. The lowest sensitivities correspond to combinations of β_1_, β_2_, and δ. For *P. aeruginosa*, the third eigenvalue is 50-fold larger than the first, and corresponds to δ. The lowest sensitivities correspond to β_1_, β_2_, and τ. For *S. aureus*, the third eigenvalue is 100-fold larger than the first, and corresponds to β_1_. The lowest sensitivities correspond to combinations of β_1_, δ, and τ. The relatively low sensitivity of τ in the LSA is consistent with the very sharp transition between the two phases, as the precise value of τ is ill-determined between the discretely sampled time points. The higher sensitivity of τ for *E. coli* is consistent with the extrapolation of the phase 2 relaxation curve to a more precise value of τ.

**Figure 4.**
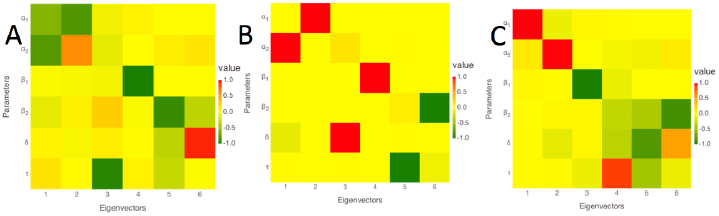
Local sensitivity analysis (LSA) results for A) *E. coli;* B) *P. aeruginosa;* and C) *S. aureus*. Columns correspond to eigenvectors, ordered left to right from the most to the least sensitive, as in Table 3. Rows correspond to parameters, with labels indicated at the left. The strength and sign of the coefficient of each parameter with each eigenvector is indicated using the color code in the legend to the right.

**Table 3.**
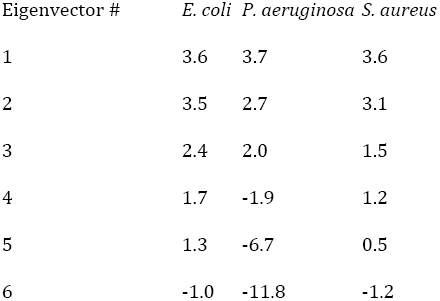
Sensitivities calculated from eigenvalues corresponding to each eigenvector from LSA. Sensitivity values are calculated as −Log_10_(λ), where λ is the eigenvalue. The connection of each eigenvector to the model parameters is visualized in Fig. 4.

## Bayesian sensitivity analysis (BSA)

We used BSA with Markov Chain Monte Carlo (MCMC) sampling (Methods) to obtain insight into probability distribution functions (PDFs) of parameter values (Figs. 5–7). Because the measurement errors were not available for individual data points, we used an arbitrary error of 1 ng/mg for each data point (Methods). The resulting PDFs therefore should not be interpreted as actual posterior PDFs, but only as indicators of: (1) the overall shape of the distributions; (2) the relative degree to which different parameters are constrained; and (3) the relative degree and nature in which different parameters are correlated.

**Figure 5.**
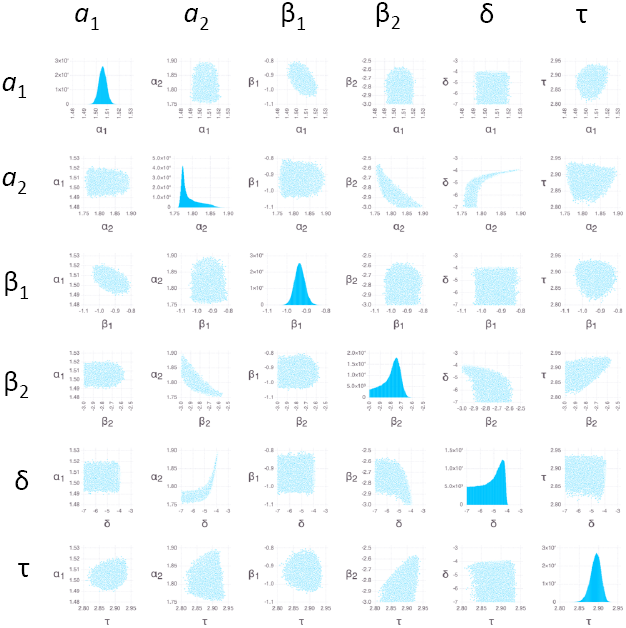
Bayesian sensitivity analysis for *E. coli*. Pairwise parameter PDFs are shown using scatter plots in off-diagonal panels; Monte Carlo sampled points illustrate the extent of the PDFs in the space of parameter values indicated on the x- and y-axes. Panels in transposed positions within the grids contain identical information with the x- and y-axes swapped. PDFs for individual parameters are shown on the diagonals using histograms, with the number of counts in each bin (out of 10^6^ total samples) indicated on the y-axis.

The parameters β_1_, β_2_, and τ in *P. aeruginosa* have relatively flat PDFs that deviate substantially from a normal distribution (Fig. 6). Sharpening the likelihood function by decreasing the errors 10-fold led to sharper PDFs for peaked distributions (Supplementary Figs. S1–S3) while maintaining similarly flat distributions for these parameters (Fig. S2), providing evidence that the flat distributions are not an artifact of the sampling.

**Figure 6.**
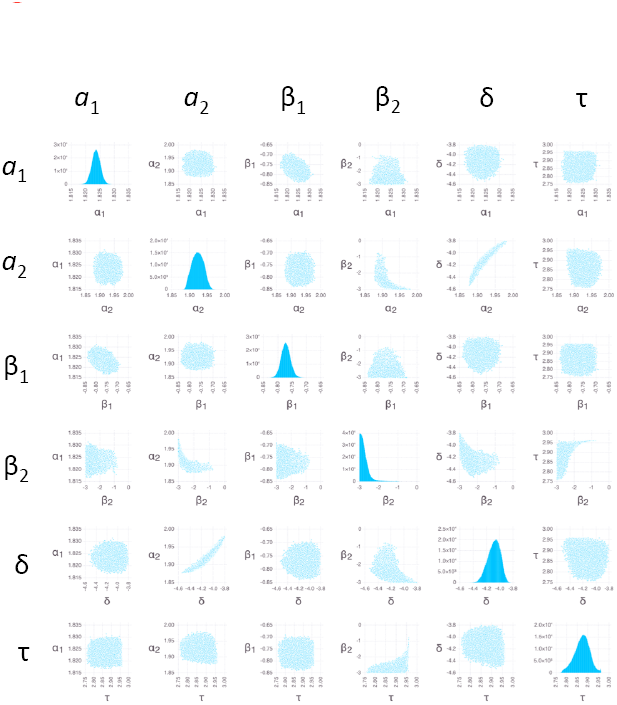
Bayesian sensitivity analysis for *P. aeruginosa*. Plots of posterior PDFs for parameter values and pairwise distributions are as in Fig. 5.

The individual PDFs for the parameters of both *E. coli* (Fig. 5) and *S. aureus* (Fig. 7) all show peaks, which is justification for the providing the estimates for all parameters in Table 1 (the distribution of β_2_ for *S. aureus* is truncated at low values due to the restricted range of the prior; however, the peak clearly falls within the range of the prior when the likelihood function is sharpened (Supplementary Figure S3)). The range of the values of β_2_ and δ among the MCMC sampled points (an indicator of the width of the PDFs) is greater than for other parameters (Table 4), indicating that estimates of β_2_ and δ are less reliable (there is a noticeable tail in the *a*_2_ distribution for *E. coli*, but the range of 0.14 Log_10_ units is still sufficiently narrow to provide an estimate). The pairwise scatter plots indicate relatively strong correlations between *a*_2_ and both β_2_ and δ. These correlations follow from Eq. (4), in which both β_2_ and δ appear in factors that multiply *a*_2_; a weaker correlation between δ and β_2_ is explained by the same term. The correlations involving δ are only substantial when δ is sufficiently large; this is because the effect of δ is negligible when it is smaller than about 10^−5^ s^−1^ (Eq. 4, considering the experiment duration of 3600 s). In addition, β_1_ appears in the term modulating *a*_1_ in Eq. (4), explaining a weak correlation between these two.

**Figure 7.**
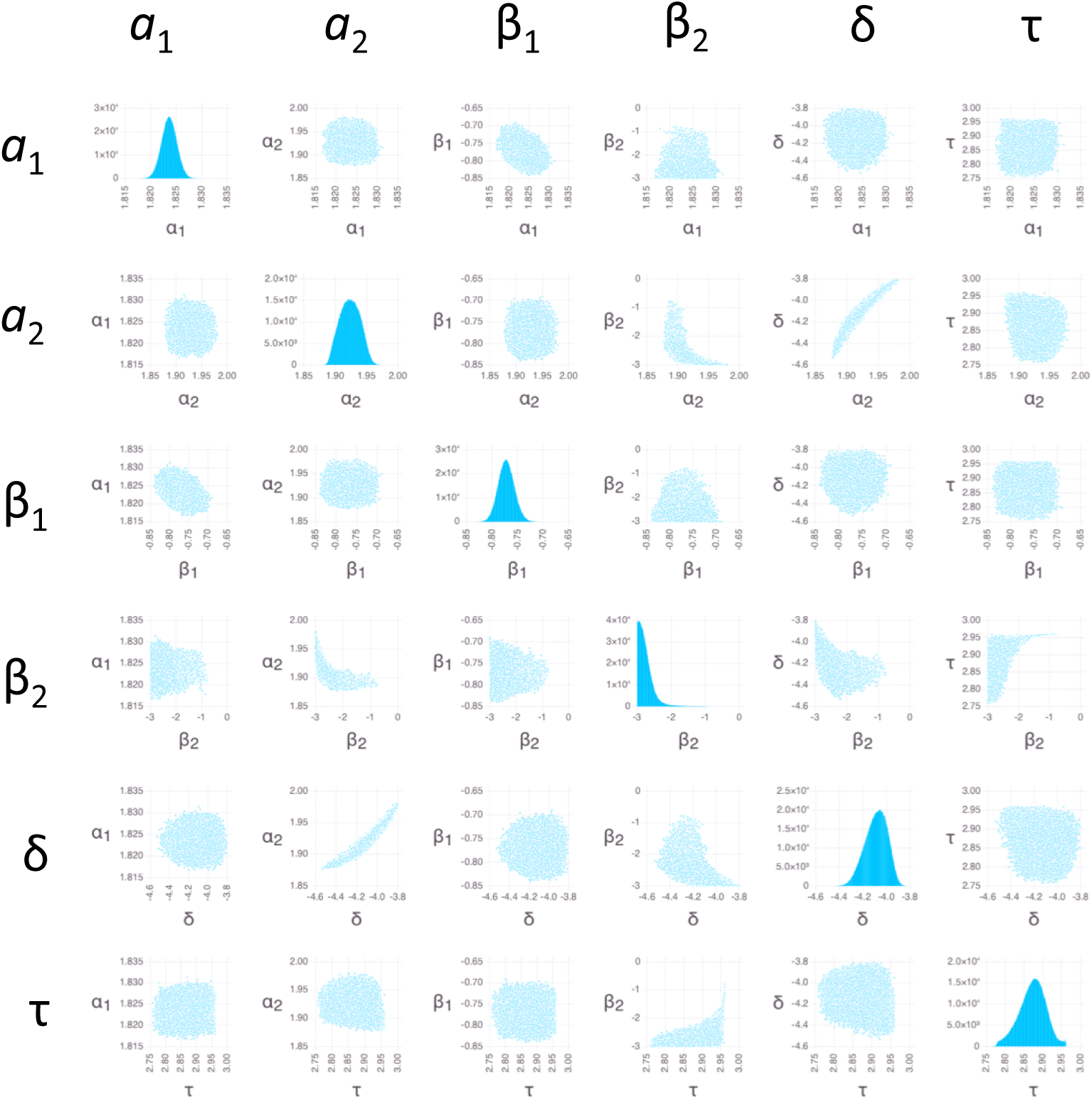
Bayesian sensitivity analysis for *S. aureus*. Plots of posterior PDFs for parameter values and pairwise distributions as in Fig. 5.

**Table 4.** Minimum and maximum values of parameters among 10^6^ random samples obtained by BSA MCMC (Log10 transformed values). Units for parameters are as in Table 1.

| | $\text{Log}_{10} a_1$ | $\text{Log}_{10} a_2$ | $\text{Log}_{10} \beta_1$ | $\text{Log}_{10} \beta_2$ | $\text{Log}_{10} \delta$ | $\text{Log}_{10} \tau$ |
| --- | --- | --- | --- | --- | --- | --- |
| *E. coli* |  |  |  |  |  |  |
| min | 1.49 | 1.75 | -1.05 | -3.00 | -7.00 | 2.81 |
| max | 1.52 | 1.90 | -0.80 | -2.57 | -3.90 | 2.94 |
| range | 0.03 | 0.14 | 0.25 | 0.43 | 3.10 | 0.13 |
| *P. aeruginosa* |  |  |  |  |  |  |
| min | 0.76 | 1.08 | -1.00 | -1.00 | -3.75 | 1.53 |
| max | 0.97 | 1.18 | 4.00 | 4.00 | -3.28 | 1.69 |
| range | 0.21 | 0.10 | 5.00 | 5.00 | 0.47 | 0.16 |
| *S. aureus* |  |  |  |  |  |  |
| min | 1.82 | 1.88 | -0.84 | -3.00 | -4.53 | 2.76 |
| max | 1.83 | 1.98 | -0.69 | -0.77 | -3.80 | 2.96 |
| range | 0.01 | 0.10 | 0.15 | 2.23 | 0.73 | 0.20 |

Like for *E. coli* and *S. aureus*, the PDFs of *a*_1_, *a*_2_, and δ for *P. aeruginosa* show peaks (Fig. 6), justifying the entry of estimates in Table 1. In addition the PDF for τ, although flat, is confined to a very narrow range, leading to an estimate in Table 1. The lack of a peak for τ can be understood by the sharp, step-like increase (Fig. 1B) combined with the discrete sampling in time; i.e., the step can be located anywhere between the time points that mark the increase, with approximately equal probability. Step-like increases in accumulation also lead to high uncertainty in both β_1_ and β_2_ (Table 4); these parameters can assume essentially any value that is large enough to enable the antibiotic to rise between the time points that mark the increase, with equal probability. Like for *E. coli* and *S. aureus*, *a*_2_ and δ are correlated for *P. aeruginosa;* however, compared to *E. coli* and *S. aureus*, increased uncertainty masks any underlying correlations involving β_1_ and β_2_.

### Predictions of permeability, net efflux rate, and accumulation factor

We used Eqs. (5)–(7) to predict the values of parameters that directly characterize permeation, net efflux, and the degree of drug accumulation. The predictions are quantified in the form of the permeability α_*i*_ in each phase *i*, the net efflux rate ε_*i*_, which is defined as the portion of the specific export rate not associated with permeation, and the accumulation factor, ϕ_*i*_, which is the fold-increase of antibiotic concentration inside the cell compared to the external concentration. A positive value of ε is associated with a net efflux and a value ϕ<1, while a negative value is associated with a net trapping of drug and a value ϕ>1. The total value of ε can be determined by a combination of individual positive or negative fluxes, potentially involving multiple mechanisms including efflux pumps and drug binding to DNA (Discussion).

The resulting predictions of *α_i_*, ε *_i_*, and ϕ*_i_* are shown in Table 5. Predictions of *α_i_* and ε *_i_* for *P. aeruginosa* are provided as lower bounds, following from the entries for β *_i_* in Table 1. The true reliability of the predictions of ϕ *_i_* depends both on the robustness of the estimation of *αi* (Sensitivity analysis results) and the uncertainty in the factor used to convert *a_0_* to ng/mg dry weight units (Methods). The reliability of the predictions of *α_i_*, ε*_i_* additionally depends on the robustness of the estimation of β*_i_*.

**Table 5.** Predicted accumulation factors (ϕ), permeabilities (α), and net efflux (ε) rates.

| | $\phi_1$ | $\phi_2$ | $\alpha_1$ (1/s) | $\alpha_2$ (1/s) | $\epsilon_1$ (1/s) | $\epsilon_2$ (1/s) |
| --- | --- | --- | --- | --- | --- | --- |
| *E. coli* | 1.09 | 2.29 | 0.126 | 0.0029 | -0.011 | -0.0016 |
| *P. aeruginosa* | 0.39 | 0.72 | >0.039<sup>a</sup> | >0.072<sup>a</sup> | >0<sup>a</sup> | >0<sup>a</sup> |
| *S. aureus* | 2.93 | 3.75 | 0.495 | 0.0053 | -0.326 | -0.0039 |

<sup>a</sup>Entries with a ‘>’ indicate a lower bound.

The predictions of permeability behavior are strikingly similar for *E. coli* and *S. aureus*. At early times, the permeability is high, corresponding to about a 5.5 s half-equilibration time for *E. coli*, and a 1.5 s half-equilibration time for *S. aureus* (α_1_ from Table 5). The permeability then decreases dramatically after about 12.5 minutes (τ from Table 1): by a factor of ̴50 for E. coli and a factor of ̴100 for S. aureus (α_1_/α_1_ from Table 5). In addition, the value of ε at both early and late times is negative for both *E. coli* and *S. aureus*, and the values of ϕ are greater than one, indicating a net trapping of antibiotic.

In contrast to *E. coli* and *S. aureus*, the values of ϕ are less than one for *P. aeruginosa*, indicating a net efflux of antibiotic. The lower bound of α_1_ for *P. aeruginosa* is consistent with *E. coli* and *S. aureus* but the lower bound of α_2_ is higher by a factor O(10). The upper bounds of ε for *P. aeruginosa* are consistent with the estimates for *E. coli* and *S. aureus*.

## Discussion

The present modeling and analysis of ofloxacin accumulation time courses indicates that *E. coli* and *S. aureus* exhibit a similar two-phase ofloxacin accumulation behavior. Whereas the two-phase behavior for *E. coli* was easily visible in the data, for *S. aureus* it was only detected as a result of the BSA. Even though the change in antibiotic level entering phase 2 is small for *S. aureus*, the BSA indicates a clear peak in the posterior distributions of all of the influx and outflux parameters, increasing confidence in the two-phase model for this case (Fig. 7).

The contribution of sensitivity analysis to the present study was substantial. The BSA enabled us to identify a significant two-phase model for *S. aureus* that we were unable to detect in the data by eye. In addition, the BSA enabled us to identify correlations in the data that are consistent with the way the parameters work together in the model. We also note that the lack of individual error estimates for each data point in the present study prevented calculation of true posterior PDFs; the availability of error estimates for future datasets will enable a substantial increase in the information provided by the BSA. The sensitivity analysis also enables a quantitative means of distinguishing of important from unimportant parameter values in the modeling. This further enables a degree of confidence to be associated with results based on the quantitative analysis, which can be used to assess which predictions are more or less likely. Some aspects of the sensitivity analysis confirm what is already apparent in just looking at the data, such as the increased importance of δ for *P. aeruginosa* compared to *E. coli;* however, even in this case, it provides quantitative measures that can enable obvious results to be obtained automatically. Automation will become increasingly important as we increase the number of data sets being analyzed. In addition, future data sets might not be as straightforward to interpret by eye, and the present study serves as an example that demonstrates the consistency of the sensitivity analysis with the interpretation of a relatively clear-cut case, so that the methods can be applied with increased confidence to cases where the interpretation might not be so obvious.

The rapid initial accumulation of ofloxacin in *E. coli, P. aeruginosa*, and *S. aureas* has been cited as evidence that these bacteria lack a substantial permeability barrier for fluoroquinolones [7]. Our modeling supports the high permeability at early times in the accumulation dynamics. The permeation rates in Table 5 lead to a half-equilibration time for ofloxacin permeation of 5.5 s into *E. coli*,, < 18 s into *P. aeruginosa*, and 1.5 s for *S. aureus*. These values are similar to the expected 5 s half-equilibration time for permeation of norfloxacin across a lipid bilayer [10]; as measured in Ref. [7], the apparent partition coefficient of ofloxacin is 4-fold greater than that of norfloxacin, so their permeabilities across the cytoplasmic membrane are expected to be within a similar range.

In contrast with the early time behavior, after about 12.5 mins, both *E. coli* and *S*. *aureus* show a predicted 50-to-100-fold decrease in permeability (Table 5). This sudden decrease in permeability only was detected using the modeling and was not detected in the prior qualitative analysis of the data [7]. One way to test this prediction is to shift the cells to a medium with radiolabeled drug, looking for a difference in labeled drug accumulation in phase 1 versus phase 2. At this point we can only speculate about the mechanism. *E. coli* is gram negative, whereas *S. aureus* is gram positive, suggesting the common decrease in permeability does not involve the outer membrane. That the permeability rates are consistent with diffusion across the cytoplasmic membrane indicates that the decrease in permeability is not necessarily be associated with transporters. It is possible that the sacculus, a mesh structure common to both *E. coli* and *S. aureus*, might be involved in a decrease in permeability. The ability of the sacculus to allow translocation of proteins as large as 25 kDa [11] or 100 kDa [12] would seem to rule out a role for the sacculus by itself, as ofloxacin has a molecular weight of 361 Da, and should ordinarily freely diffuse across the mesh. However, the sacculus is associated with a comparable mass of other biomolecules in both gram negative and positive bacteria [13] [14]. If ̴98% of the mesh were to become blocked somehow, it could explain the ̴50-fold decrease in permeability for *E. coli*.

For *E. coli* and *S. aureus*, the values of the accumulation factor ϕ>1 indicate net trapping of antibiotic. The trapping is weak for *E. coli* in phase 1 (ϕ_1_=1.09) and becomes stronger in phase 2 (ϕ_2_=2.29). Compared to *E. coli*, the trapping is stronger in *S. aureus* for both phases: ϕ_1_=2.93 and ϕ_2_=3.75. For *P. aeruginosa*, the values of ϕ 1 indicate a net efflux of antibiotic that is stronger in phase 1 (ϕ_1_=0.39) than in phase 2 (ϕ_2_=0.72).

Several individual effects can combine to determine whether there is a net trapping or efflux of antibiotic. Targeted experiments will be required to determine the role of specific mechanisms, but we can gain some insights by assuming that an important effect favoring trapping is the immobilization of antibiotic on DNA. This effect can be estimated using the dissociation constant *K_d_* = 633 μM reported by Shen et al. [15] for nonspecific binding to DNA of norfloxacin at high concentrations (10–1000 μM). Assuming roughly 2.1 genome equivalents per cell [16], a volume of 1.25 fL, and 4.7 × 10^6^ bp per chromosome yields an estimated DNA bp concentration in *E. coli* [D] = 1.3 × 10^4^ μM. Then, assuming the *K_d_* for norfloxacin applies to ofloxacin and that each bp can bind one drug molecule yields a fraction of DNA-bound drug of 1/(1+*K_d_*/[D]) = 0.95, or a mobile fraction of just 0.05. In this case, the contribution of permeability to the outflow would be only 5% of that assumed in Eq. (6), leading to an increase in efflux by 0.95α. This translates into an increase in the specific efflux rate by 0.12 s^−1^ for *E. coli* in phase 1.

Based on the above estimates, despite the small negative value of the net efflux parameter ε_1_ (= −0.011 s^−1^) in Table 5, due to the effect of trapping by DNA binding, we expect the underlying rate of efflux in *E. coli* to be 0.11 s^−1^ (= 0.12 s^−1^−0.011 s^−1^), which is comparable to the rate of permeation. To assess the importance of efflux in *S. aureus*, we use similar reasoning as for *E. coli*, noting that the net efflux parameter ε_1_ is −0.326 s^−1^ and the effective permeability α is 0.495 s^−1^ (Table 5). Assuming the same degree of trapping in *S. aureus* as in *E. coli* yields a contribution 0.48 s^−1^ and an estimated specific efflux rate of roughly 0.15 s^−1^ (= 0.48 s^−1^ − 0.326 s^−1^) in phase 1, which is close to the value for *E. coli*. In contrast to *E. coli*, however, where the efflux rate is comparable to the inflow rate, in *S. aureus* the efflux rate is less than 1/3 of the inflow rate.

The estimated specific efflux rates in phase 1 of 0.11 s^−1^ for *E. coli* and 0.15 s^−1^ for *S. aureus* can be used to estimate the rate of outflow of drug due to efflux. Multiplying by *a*_1_ yields an outflow of 3.5 ng/mg/s (= 0.11 s^−1^ × 32 ng/mg) for *E. coli*, and 10 ng/mg/s (= 0.15 s^−1^ × 66.6 ng/mg) for *S. aureus*. Dividing by the molecular weight of ofloxacin (361.368 g/mol) yields 0.0097 nmol/mg/s for *E. coli* and 0.028 nmol/mg/s for *S. aureus*. These values are similar to the *Vmax* value of 0.0235 ± 0.003 nmol/mg/s found by Lim and Nikaido [6] for efflux of nitrocefin from *E. coli*, and are much lower than the value of 1.82 ± 0.85 nmol/mg/s found for cephaloridine efflux from *E. coli* in the same study. Therefore our predictions are consistent with the *V_max_* for efflux of ofloxacin in *E. coli* and *S. aureus* being similar to the *Vmax* for efflux of nitrocefin in *E. coli*.

In addition to DNA binding, differential permeability of ionic subspecies of ofloxacin or Mg2+ chelates of ofloxacin might also influence the balance of inflows and outflows [10]. These effects can easily lead to two-fold or greater accumulation or exclusion of antibiotic [10]. In addition to the predicted change in membrane permeability, therefore, changes in DNA binding, efflux, pH, and the availability of Mg2+ all might potentially be linked to the two-phase behavior of ofloxacin accumulation.

Because we predict that the underlying efflux rate is comparable to the inflow rate in *E. coli*, we also predict that efflux plays an important role in ofloxacin accumulation. In contrast, for *S. aureus*, we predict efflux plays a less important role in ofloxacin accumulation, because the efflux rate is predicted to be substantially lower than the inflow rate. For *P. aeruginosa*, we are unable to make predictions based on quantitative estimates of rate constants, due to uncertainty in the key parameter values in Table 1 and Table 5. However, because the internal concentration of antibiotic is predicted to be less than the external concentration (ϕ < 1 in Table 5), we predict that efflux is important in *P. aeruginosa* accumulation of ofloxacin.

Both *P. aeruginosa* and *S. aureus* exhibit a visible slow decrease in accumulated drug during the second phase (Fig. 1). The model captures this decrease in terms of the parameter δ which increases β_2_ by a factor [1 + δ(t—τ)] in Eq. (3). Mechanisms for increasing β include enhancing efflux (e.g., by increased expression of pumps) or decreasing trapping (e.g., by chromosome remodeling that results in a decrease in drug binding to DNA). Importantly, the combination of Eq. (4) and Eq. (5) indicate that 6 can be interpreted alternatively as decreasing012 by a factor [1 + δ(t—τ)]^−1^. Thus, in addition to an increase in efflux or a decrease in trapping, the acclimation behavior can be attributed to a decrease in permeability (e.g., through the hypothetical mechanisms described above). Additional experiments targeted at the mechanism of decrease will be required to determine the cause of the acclimation response.

Ofloxacin is bactericidal at the concentration of 10 mg/L used in the experiments [7], which might have important implications for the interpretation of the results. If nonviable cells are excluded from the analyzed pellet after centrifugation (e.g. because they have been lysed and remain suspended), the accumulation data simply correspond to viable cells, and the interpretation is straightforward. If nonviable cells are included in the pellet, the interpretation depends on the relative populations of viable and nonviable cells. According to Ref. [7], 5 mins after exposure in phosphate buffer, the viable count of *E. coli* decreased by 0.74 Log10 units, the count of *P. aeruginosa* decreased by 0.45 Log_10_ units, and the count of *S. aureus* decreased by 0.4 Log_10_ units. These data indicate that in all cases, the number of viable cells decreases by at least 2.5-fold after 5 mins. At the longest times, 60 mins after exposure, the values were 2.06, 3.75, and 0.54 Log_10_ units, respectively, indicating that the number of viable cells of *E. coli* and *P. aeruginosa* decreases by at least 100-fold over the course of the experiment. These results indicate that if nonviable cells are included in the analyzed pellet, the behavior should be dominated by the nonviable cells; indeed, trapping of antibiotic in nonviable cells removes antibiotic from the environment and can increase the chance of survival of neighboring viable cells, leading to a fitness advantage at the population level.

That the model matches the data well (Fig. 1) is consistent with the interpretation that the behavior is predominantly due to one type of cell, either viable or nonviable. However, the killing kinetics do indicate that the number of viable cells is still not negligible at early times for all cells, and at late times for *S. aureus*. In addition, including the effect of antibiotic on cell growth and death will be critical to connecting the present results to drug resistance. Future work should therefore include investigating an expanded dynamical model that includes subpopulations of viable and nonviable cells, including the dependence of cell growth (for bacteriostatic effect) and death (for bactericidal effect) on intracellular antibiotic concentration. Including the antibiotic effect (e.g., via the target inhibitory concentration [4–6]) is essential for understanding the relation between the amount of accumulated drug and antibacterial activity, as they are not necessarily correlated [7]. In particular, Nagano and Nikaido [5] used mathematical modeling to predict MIC values of drugs for *E. coli*, and to show how efflux pump deletion or overexpression might not change the MIC value of drugs that are nevertheless strong efflux pump substrates.

It is interesting to consider the importance of models of drug accumulation in light of a recent survey of physical properties of active compounds in a drug screening collection and their relation to whole cell antibacterial activity [17]. The survey reported on the difficulties encountered in simultaneously optimizing both biochemical potency and antibacterial activity, and concluded that “what is clearly needed is greater insight into medicinal chemistry strategies which optimize transport through porins and decrease efflux through the prolific efflux pumps.” Together, the findings reported here and elsewhere [5,6] suggest that mathematical modeling of drug accumulation can provide these insights, and thus can be a key tool in enabling antibacterial medicinal chemistry.

## Materials and Methods

### Model

Our dynamical model of drug accumulation lumps together transport across the inner and outer membranes. This is appropriate as the drug target is in the cytoplasm, there are no data on drug accumulation in the periplasm, and we wish to use the same model for gram positive or negative bacteria. We note that this lumping does not prevent the interpretation of results in light of differences between the outer and inner membranes (Discussion).

The time dependence of accumulated drug *a(t)* is modeled using

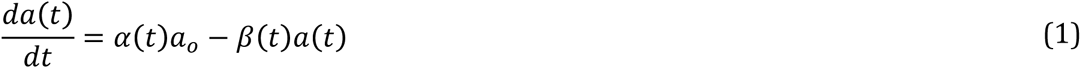

where *a_o_* is the environmental antibiotic concentration, α(t) is the specific rate of increase, and β(t) is the specific rate of decrease. The term “specific” is used to indicate that the rate constant is multiplied by the antibiotic concentration. We assume α(t) corresponds to the permeability, although it also can include active transport. We assume β(t) corresponds to both active and passive transport, although it also can include degradation and cell growth. Cell growth in *E. coli* occurs by a constant cell size extension [18], which is consistent with including cell growth in β. If included, assuming a 42 min doubling time, cell growth would contribute a specific rate of 2.75 × 10^−4^ s^−1^; this is orders of magnitude smaller than the values of β at early times (Table 1), when growth is expected to be highest, and is at least 4-fold smaller than the values of β at early times (Table 1), when growth is expected to be lowest. Ignoring cell growth therefore is a reasonable approximation in the present study, although it might be necessary to include it in other studies.

Equation (1) was developed assuming that only viable cells are analyzed. This corresponds to the case when nonviable cells are disrupted and are excluded from pellet after centrifugation in Ref. [7]. Even when nonviable cells are not excluded from analysis (Discussion), Eq. (1) applies to viable cells when the external antibiotic level is low and there is a negligible fraction of nonviable cells at all times. If nonviable cells are included in the pellet and there is a negligible fraction of viable cells at all times, Eq. (1) will still be relevant but will instead describe accumulation in the nonviable cells. If both viable and nonviable cells are analyzed and there is an intermediate drug concentration, the model should be extended to include both viable and nonviable cells, with population dynamics (Discussion).

To model the data in Fig. (1), we assume α(t) and β(t) both switch rapidly from a constant value in phase 1 to a different constant value in phase 2. The import rate α(t) is given by

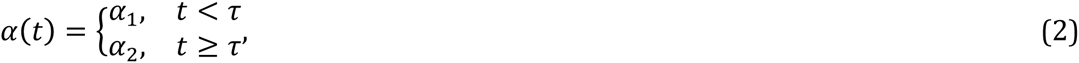

where α_1_ applies to the early phase of the accumulation dynamics (t<τ), and α_2_ applies to the late phase (t≥τ).

The export rate β(t) is given by

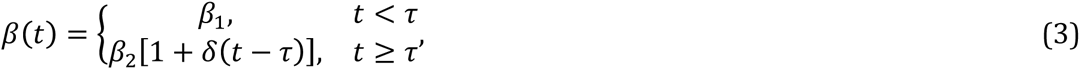

where β_1_ applies during the early phase (t<τ), β_2_ applies during the late phase (t≥τ), and δ is the fractional rate of increase of export during the later phase.

Assuming rapid relaxation (P2) compared to acclimation (6) in phase 2 yields

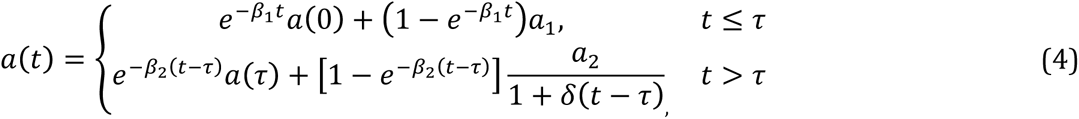

where *a*_1_ = α_1_ *a_0_* β_1_, and *a*_2_ = α_2_ *a_0_*β_2_.

Given additional assumptions, the α and β parameters can be used to derive permeation and net efflux rates (see below). The δ parameter models long-term acclimation by, e.g., gene regulation.

### Parameter estimation

Parameter estimation for Eq. (4) was performed using published experimental time courses of ofloxacin accumulation in *E. coli, P. aeruginosa*, and *S. aureus* [7]. Data points were extracted from Figure 1 in Ref. [7]. Numerical solutions to Eq. (4) were obtained for each data point using functions defined in Mathematica Version 10 (Wolfram Research, Inc., Champaign, IL). Errors for each individual data point were not available, and were therefore estimated using the maximum values given for norfloxacin in Ref. [7]: ±12.5, ±11.6, and ±9.7 ng/mg for *E. coli, P. aeruginosa*, and *S. aureus*, respectively. To obtain error estimates for ofloxacin, these maximal errors were scaled using the initial rapid phase concentration entries in Tables II, III, and IV of Ref. [7], and were then divided by two. The following were the resulting estimated errors for each data point: 2.3 ng/mg for *E. coli*, 0.75 ng/mg for *P. aeruginosa;* and 4.35 ng/mg for *S. aureus*. We also performed the fits using the maximal errors as the estimates, without dividing by two; however, we favored using the halfQmaximal errors for the following reasons: (1) the maximal values are very likely overestimates; and (2) the halfQmaximal values yielded χ^2^/NDF values close to 1, whereas the maximal values yielded values substantially lower than 1. The choice of errors between these cases merely resulted in a different overall scaling χ*^2^* and did not influence the optimal parameter values obtained in the fits.

A standard χ^2^/NDF was used as a goodness of fit measure between the model and the data, using a number of degrees of freedom NDF = number of data points minus six parameters. The χ^2^/NDF was minimized with respect to values of *a*_1_, *a*_2_, β_1_, β_2_, δ, and τ using the `FindMinimum` method, assuming a(0)=0. The values of *a*_1_, *a*_2_, β_1_, β_2_, δ were varied over twelve decades. For *E. coli* and *P. aeruginosa*, the value of τ was varied from 0–10,000 s (covering the full span of each 3,600 s time course) to seek an optimal solution. For *S. aureus*, optimization was initially performed for τ = 10,000 s (a time beyond the last data point), as the behavior appeared to be one phase. A more thorough investigation of the parameter space using Bayesian sensitivity analysis (BSA), however, revealed a relatively weak but significant two-phase response, with a value of τ that is similar to *E. coli*. Therefore the optimization for *S. aureus* also was performed varying τ from 0–10,000 s. Mathematica workbooks and experimental data for all cases are available in a compressed archive in the Supporting Information.

### Sensitivity analysis methods

All sensitivity analyses were performed using the open source code Model-Analysis and Decision Support (MADS), developed at Los Alamos National Laboratory (mads.lanl.gov).

*Global Sensitivity Analysis (GSA)*: We performed variance-based GSA [19], which decomposes the variance of the output into parts ascribed to different input parameters using a Fourier Haar decomposition [20,21]. Both the “ main effect“ and the “total effect” results presented here for each input parameter were calculated using the Sobol Monte Carlo algorithm [22] using about 10^6^ independent model evaluations drawn from log-uniform distributions defined by Table 2. We found that this number of samples was sufficient to achieve convergence for the estimated quantities. The ranges in Table 2 were chosen arbitrarily to encompass the optima in Table 1 while exploring a reasonably broad region of parameter space and satisfying the requirement of rapid relaxation (β2) compared to acclimation (δ) in phase 2 for Eq. (4).

*Local Sensitivity Analysis (LSA)*: LSA has a rich history of use in assessing the robustness of biochemical network model behavior to changes in parameter values [23,24]. In LSA, the local gradients of the model output are calculated with respect to model parameters at a fixed point (the point with the lowest χ^2^). Here we use the gradients to define a covariance matrix of variations [25]. We used a finite difference method requiring a limited number of model evaluations (equal to the number of unknown parameters). As a result, we obtain a gradient matrix **J** (i.e. the Jacobian) with dimensions [*m* × *n*] where *m* is the number of model parameters and *n* is the number of model inputs (we use logarithmic derivatives, to obtain an analysis in terms of dimensionless parameters). Each component of the **J** matrix represents the local sensitivity of each model parameter to each model output [25]. We analyze the covariance matrix **C** of model parameters which is computed as **C** = [J^T^J]^−1^ The covariance matrix is analyzed using eigenanalysis where eigenvectors and eigenvalues of the covariance matrix are explored. How dominant (important) is each eigenvector depends on the respective eigenvalues; the smaller the eigenvalue, the higher the importance of the eigenvector. The components of each eigenvector represent the contributions of each model parameter to the simultaneous variation of multiple parameters: the larger the absolute value of the components, the larger the contribution. Model parameters with a large contribution in dominant eigenvectors are important (sensitive) model parameters. Model parameters with large contribution in nonQdominant eigenvectors are unimportant (insensitive) model parameters. If several model parameters have an important contribution in a single eigenvector, these model parameters are correlated. If these contributions have the same sign, the correlation is positive. If these contributions have opposite signs, the correlation is negative.

*Bayesian Sensitivity Analysis (BSA)*: Posterior probability density functions (PDFs) of the model parameters given observed data (Figs. 5–7) were obtained following Bayes theorem [26,27]. The PDFs were obtained using a log-uniform prior with the ranges indicated in Table 2, with a likelihood function defined as *e̴*^*x*^ *I*^*2*^, using an arbitrary error value of 1 ng/mg for each data point. The error was decreased to 0.1 ng/mgto demonstrate the effect of sharpening the likelihood function in Supplementary Figs. S1–S3. The BSA was performed using the Robust Adaptive Metropolis Markov Chain Monte Carlo (MCMC) algorithm [28]. Analysis was performed using sample sizes of 10^4^,10^5^, and 10^6^; we found a sample size of 10^6^ yielded adequate sampling. The posterior PDF scatter plots (off-diagonal panels in Figs. 5–7) show an overlay of individual counts; the extent of the plots reflects the parameter range over which 10^6^ random samples can fall. The posterior parameter ranges in Table 4 were obtained by identifying the minimum and maximum parameter values among 10^6^ points sampled using the MCMC. (The values in Table 4 must be interpreted in light of the fact that the range might increase with increasing number of samples, and that the precise values can change depending on differences in sampling, e.g., using different random number seeds.) The histograms (diagonal panels in Figs. 5–7) indicate the shape of the individual marginalized PDFs using the number of counts on the y-axis.

### Predictions of permeability, net efflux rate, and accumulation factor

Estimation of the model parameters enables prediction of permeability and effluxrates. Given *a*_0_, the specific import rate α_*i*_ in each phase *i* may be calculated as

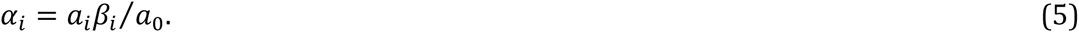

Assuming the import is due to permeation, α*_i_* is the permeability. Next, assume the specific export rate β is a sum of contributions from permeability, α and other effects. The other effects are then captured by the difference ε between β and α:

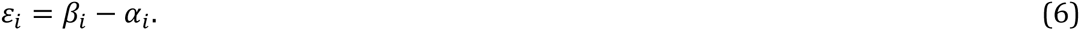

When ε_i_>0, outward flow is enhanced compared to permeation, and ε_i_ indicates a net efflux. When ε_i_<0, the outward flow is decreased compared to permeation, and ε_i_ indicates a net trapping of drug.

Finally, the accumulation factor ϕ is used as a measure of the degree to which antibiotic is either accumulated in or expelled from the cell. It is given by

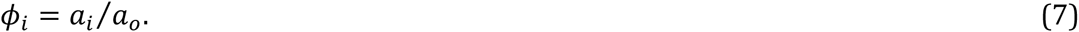

If ϕ_*i*_<1 the internal antibiotic concentration is less than the external antibiotic, which is associated with net efflux (ε_i_0). If ϕ_*i*_>1 the internal antibiotic exceeds the external antibiotic, which is associated with net trapping (ε_i_>0).

Eqs. (5)–(7) are only valid under the same assumptions as for Eq. (4). Predictions of α_*i*_and ε_i_ are only meaningful using reliable estimates of both α_*i*_ and β_*i*_. In contrast, ϕ_*i*_ can be predicted just with estimates of *a*_*i*_, and *a*_0_.

The predictions use the environmental drug concentration *a*0 in ng drug/mg dry weight units. The external ofloxacin concentration in the experiments was 10 mg/L [7]. To convert to ng/mg dry weight units, following Ref. [29], we used a buoyant density of 1.1 g/mL and a 31% dry weight for *E. coli*, a buoyant density of 1.2 g/mL and a 48% dry weight for *P. aeruginosa* (transferred from *P. putida*), and assumed a buoyant density of 1.1 g/mL and a 40% dry weight for *S. aureus*. This yielded conversion factors of 0.341 (mg drug/L)/(ng drug/mg dry weight) for *E. coli*, 0.528 for *P. aeruginosa*, and 0.440 for *S. aureus*. The uncertainty in the dry weight fraction for *S. aureus* should be estimated at about 25%, given the range spanned by *E. coli* and *P. aeruginosa*. This introduces uncertainty in the predictions beyond those indicated by the sensitivity analysis alone.

## Acknowledgments

We are grateful to the reviewers for their comments, which led to substantial improvements in the paper. This study was performed under the auspices of the LaboratoryQDirected Research and Development Program at Los Alamos National Laboratory, which is managed for the US Department of Energy by Los Alamos National Security, LLC under contract DEQAC52Q06NA25396.

---

**Figure S1.**
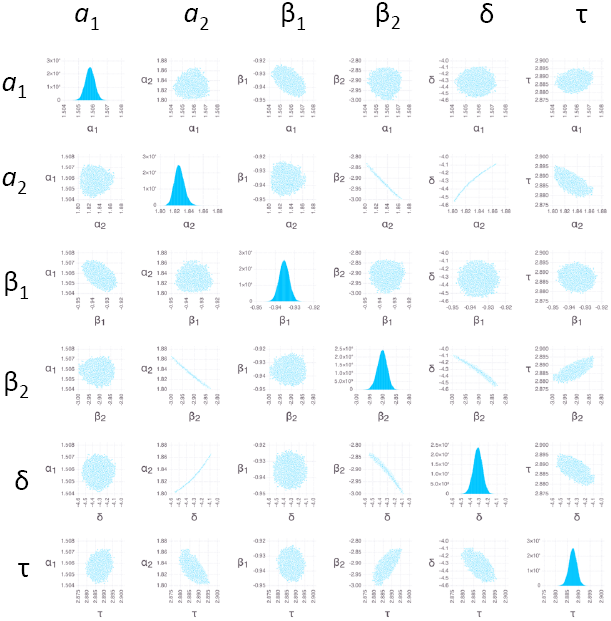
Bayesian sensitivity analysis for *E. coli* using a sharpened likelihood function. The analysis and presentation is the same as in Fig. 5, but using a 10-fold lower error value.

**Figure S2.**
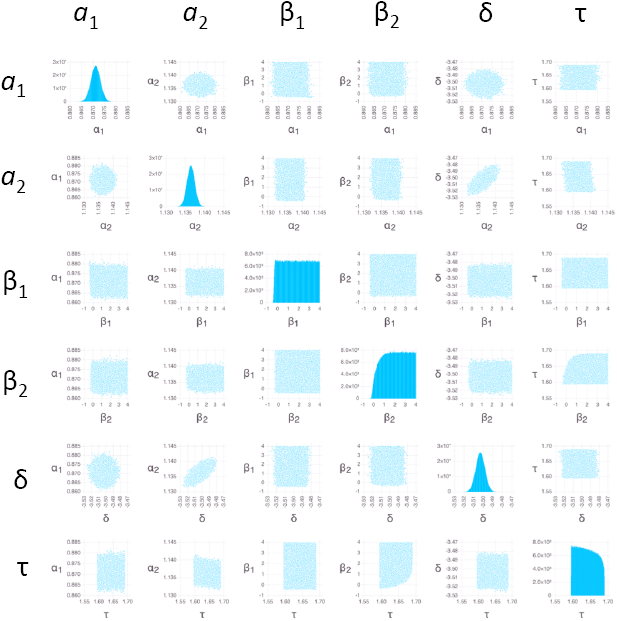
Bayesian sensitivity analysis for *P. aeruginosa* using a sharpened likelihood function. The analysis and presentation is the same as in Fig. 6, but using a 10-fold lower error value.

**Figure S3.**
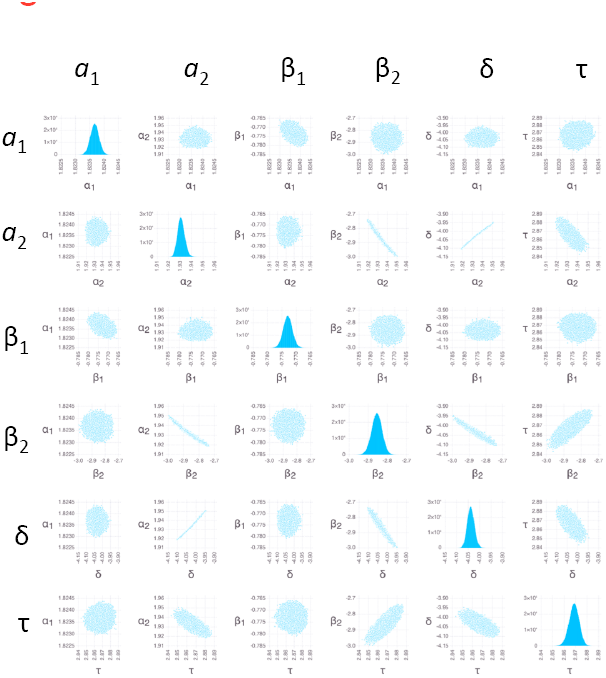
Bayesian sensitivity analysis for *S. aureus* using a sharpened likelihood function. The analysis and presentation is the same as in Fig. 7, but using a 10-fold lower error value.

## References

1. Li XZ, Nikaido H (2009) Efflux-mediated drug resistance in bacteria: an update. Drugs 69: 1555–1623.

2. Kourtesi C, Ball AR, Huang YY, Jachak SM, Vera DM, et al. (2013) Microbial efflux systems and inhibitors: approaches to drug discovery and the challenge of clinical implementation. Open Microbiol J 7: 34–52.

3. Nikaido H, Pages JM (2012) Broad-specificity efflux pumps and their role in multidrug resistance of Gram-negative bacteria. FEMS Microbiol Rev 36: 340–363.

4. Nikaido H, Normark S (1987) Sensitivity of Escherichia coli to various beta- lactams is determined by the interplay ofouter membrane permeability and degradation by periplasmic beta-lactamases: a quantitative predictivetreatment. Mol Microbiol 1: 29–36.

5. Nagano K, Nikaido H (2009) Kinetic behavior of the major multidrug efflux pump AcrB of Escherichia coli. Proc Natl Acad Sci U S A 106: 5854–5858.

6. Lim SP, Nikaido H (2010) Kinetic parameters of efflux of penicillins by the multidrug efflux transporter AcrAB-TolC of Escherichia coli. Antimicrob Agents Chemother 54: 1800–1806.

7. Asuquo AE, Piddock LJ (1993) Accumulation and killing kinetics of fifteen quinolones for Escherichia coli, Staphylococcus aureus and Pseudomonas aeruginosa. J Antimicrob Chemother 31: 865–880.

8. Diver JM, Piddock LJ,Wise R (1990) The accumulation of five quinolone antibacterial agents by Escherichia coli. J Antimicrob Chemother 25: 319–333.

9. Thanassi DG, Suh GS, Nikaido H (1995) Role of outer membrane barrier in efflux- mediated tetracycline resistance of Escheric2hia coli. J Bacteriol 177: 998–1007.

10. Nikaido H, Thanassi DG (1993) Penetration of lipophilic agents with multiple protonation sites into bacterial cells: tetracyclines and fluoroquinolones as examples. Antimicrob Agents Chemother 37: 1393–1399.

11. Demchick P, Koch AL (1996) The permeability of the wall fabric of Escherichia coli and Bacillus subtilis. J Bacteriol 178: 768–773.

12. Vazquez-Laslop N, Lee H, Hu R, Neyfakh AA (2001) Molecular sieve mechanism of selective release of cytoplasmic proteins by osmotically shocked Escherichia coli. J Bacteriol 183: 2399–2404.

13. Brown S, Santa Maria JP Jr., Walker S (2013) Wall teichoic acids of gram-positive bacteria. Annu Rev Microbiol 67: 313–336.

14. Silhavy TJ, Kahne D, Walker S (2010) The bacterial cell envelope. Cold Spring Harb Perspect Biol 2: a000414.

15. Shen LL, Baranowski J, Pernet AG (1989) Mechanism of inhibition of DNA gyrase by quinolone antibacterials: specificity and cooperativity of drug binding to DNA. Biochemistry 28: 3879–3885.

16. Neidhardt FC, Umbarger HE (1996) Chemical Composition of *Escherichia coli*. In: Neidhardt FC, Curtiss III R, Ingraham JL, Lin ECC, Low KB et al., editors. Escherichia coli and Salmonella: Cellular and Molecular Biology. 2 ed. Washington, D. C.: ASM Press.

17. Brown DG, May-Dracka TL, Gagnon MM, Tommasi R (2014) Trends and exceptions of physical properties on antibacterial activity for Gram-positive and Gram-negative pathogens. J Med Chem 57: 10144–10161.

18. Campos M, Surovtsev IV, Kato S, Paintdakhi A, Beltran B, et al. (2014) A constant size extension drives bacterial cell size homeostasis. Cell 159: 1433–1446.

19. Sobol' IM (1990) On sensitivity estimation for nonlinear mathematical models. Matematicheskoe Modelirovanie 2: 112–118.

20. Sobol' IM (1993) Asymmetric convergence of approximations of the Monte Carlo method. Computational mathematics and mathematical physics 33: 1391–1396.

21. Saltelli A, Andres TH, Homma T (1993) Sensitivity Analysis of Model Output - an Investigation of New Techniques. Computational Statistics & Data Analysis 15:211–238.

22. Saltelli A, Ratto M, Andres T, Campolongo F, Cariboni J, et al. (2008) Global Sensitivity Analysis: The Primer: John Wiley & Sons.

23. Savageau MA (1976) Biochemical systems analysis: a study of function and design in molecular biology. Reading, MA: Addison-Wesley.

24. Savageau MA (2013) System design principles. In: Wall ME, editor. Quantitative Biology: From Molecular to Cellular Systems. Boca Raton: Taylor and Francis. pp. 23–49.

25. Gustafson P, Srinivasan C, Wasserman L (1995) Local sensitivity analysis. Bayesian Statistics 5: 631–640.

26. Oakley JE, O'Hagan A (2004) Probabilistic sensitivity analysis of complex models: a Bayesian approach. J Roy Stat Soc B 66: 751–769.

27. Weiss R (1996) An approach to Bayesian sensitivity analysis. J Roy Stat Soc B 58: 739–750.

28. Vihola M (2012) Robist adaptice Metropolis algorithm with coerced acceptance rate. StatComp 22.

29. Bratbak G, Dundas I(1984) Bacterial dry matter content and biomass estimations. Appl Environ Microbiol 48: 755–757.

